# INLAomics for Scalable and Interpretable Spatial Multiomic Data Integration

**DOI:** 10.1101/2025.05.02.651831

**Authors:** Lukas Arnroth, Sanja Vickovic

## Abstract

Integrating spatial transcriptomics with antibody-based proteomics enables the investigation of biological regulation within intact tissue architecture. However, current approaches for spatial multi-omics integration often depend on dimensionality reduction or autoencoders, which disregard spatial context, limit interpretability, and face challenges with scalability. To address these limitations, we developed INLAomics, a multivariate hierarchical Bayesian framework that models protein abundance in tissue sections by leveraging histological features and latent spatial factors inferred from spatial transcriptomics data. INLAomics supports two key applications: (1) identifying spatial gene co-expression programs to build interpretable gene–protein networks, and (2) predicting spatial protein expression in tissues lacking proteomics measurements. Applied across diverse datasets, INLAomics reveals previously unrecognized gene–protein associations and achieves substantial improvements in protein prediction accuracy over models that treat each modality independently. The framework is both computationally efficient and biologically interpretable, offering a scalable solution for integrative analysis of spatial multi-omics data

## 2 Introduction

The functionality of cells within multicellular systems hinges on intricate interactions and division of labor within tissue architectures, collectively forming dynamic organ-wide ecosystems (Palla et al., 2022). Disruptions to these ecosystems, such as those occurring in disease states, can unveil crucial insights into the correlation between tissue structure and its phenotypic function. Understanding these structural and molecular alterations holds promise for deciphering disease mechanisms (Hanahan and Coussens, 2012). Recent innovations have enabled the capture of high-dimensional measurements from single cells within their native spatial context, facilitating a more comprehensive understanding of cellular dynamics (Ståhl et al., 2016; Vickovic et al., 2019; Ke et al., 2013; Lee et al., 2014; Lubeck et al., 2014; Chen et al., 2015; Codeluppi et al., 2018; Moffitt et al., 2018; Rodriques et al., 2019). Some of these approaches have also been adapted to study both mRNA expression and antibody-antigen interactions simultaneously and at scale (Stoeckius et al., 2017; Vickovic et al., 2022; Ben-Chetrit et al., 2023; Liu et al., 2023) (Fig. 1a, b) facilitating the study of spatially co-regulated biological processes. However, as gene expression undergoes regulation across various stages, from transcription to protein degradation (Rabani et al., 2014; Jovanovic et al., 2015), the correlation between protein concentrations and RNA abundances has often been reported as weak due to various protein regulatory pathways (Fangma et al., 2023). Previous studies have shown that integrating analysis from multiple layers of molecular information can identify subtle variations and interactions that may be missed by analyzing each data type independently (Nativio et al., 2020). However, computational methods that address spatial multi-modal variation from large tissue cohorts for multi-parameter integration are lacking.

**Fig. 1:**
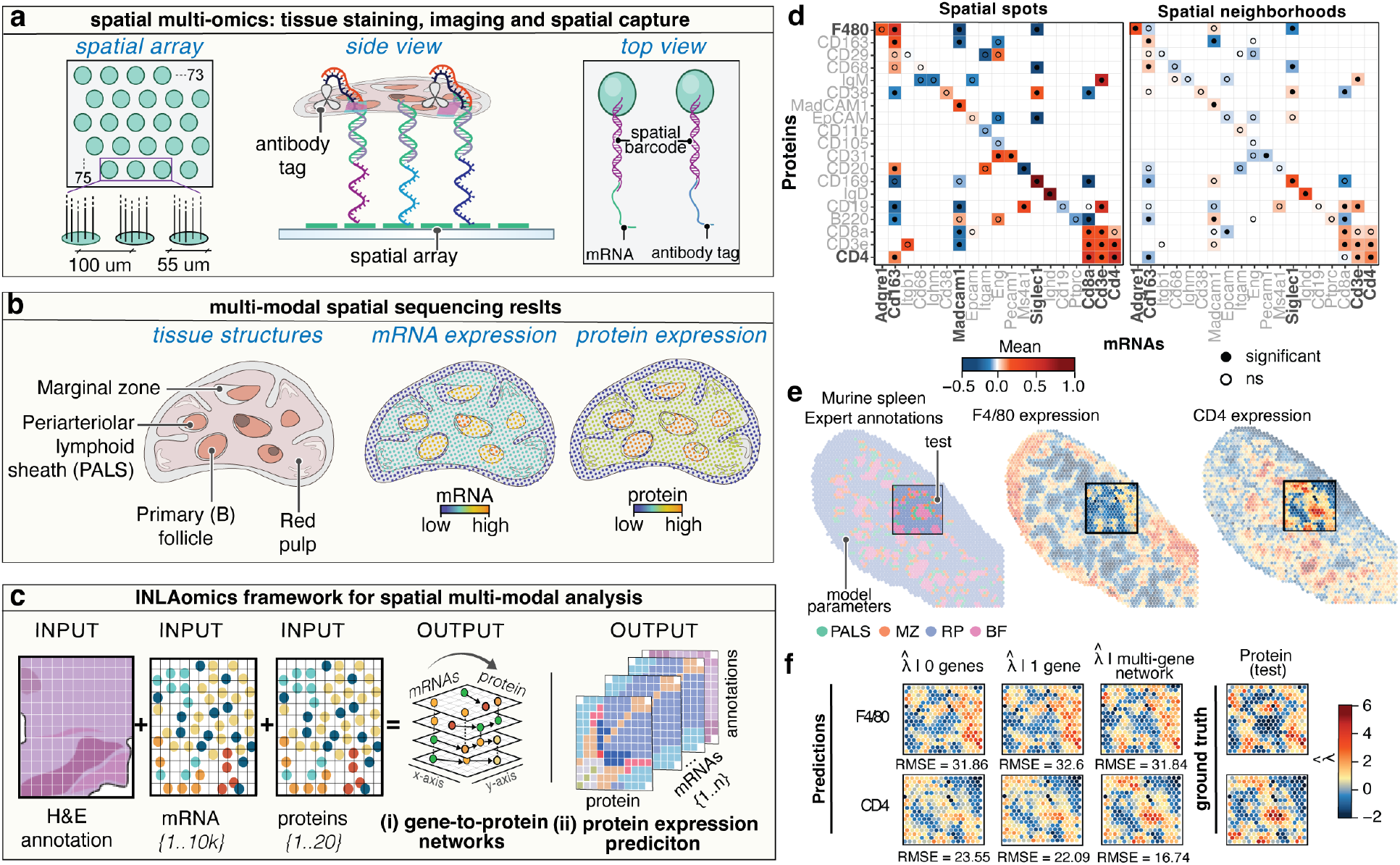
INLAomics for spatial multi-omics data integration, **a-c**, Spatial multi-omics data generation **(a)**, illustrative results when applied to murine spleen tissues **(b)** and description of the INLAomics workflow **(c). d**, Visualization of a gene-to-protein tissue network constructed via sequential model selection (Suppl. Tab. 2) for proteins and gene pairs showing cross-assay parameter estimates (color) of spot-to-spot (left) and neighbors-to-spot (right) effects (ns = non significant). **e**, Spatial multiomics murine spleen data visualized trough expert annotations (color code) and protein expression of F4/80 and CD4 (color scale). Test inset marked (black rectangle). **f**, Prediction of F4/80 and CD4 protein insets in spatial multi-omics murine spleen data based on gene-protein pairs and the gene-protein networks determined from 200 most highly expressed genes which when added sequentially to the model according to their association strength, improved model predictions. Three protein models (left)— unconditional (0 genes), paired (protein conditional on gene, 1 gene), and conditional multi-gene network (protein conditional on gene pair and any number of highly expressed genes)— are compared by RMSE and visually (color scale) to ground truth protein expression (right).

Existing methods primarily rely on dimensionality reduction, a key component of many single cell multimodal data integration approaches, or factor analysis and canonical correlations to project data onto a low dimensional space (Velten et al., 2022; Brown et al., 2023; Jiang et al., 2023). Nonetheless, such approaches tend to ignore the spatial coordinates that are the distinguishing feature of spatial multi-omics data. In other cases, transformed counts are approximated as normally distributed, and subsequently modeled as Gaussian random fields (Sun et al., 2020; Weber et al., 2023). Such approaches suffer from approximation errors in distributions and might not be computationally feasible for estimating large multi-modal datasets. Several deep learning methods have also been developed, with variational auto encoders (VAEs) commonly used to embed multi-modal data into a shared latent space (Stark et al., 2020; Gong et al., 2021; Poirion et al., 2021; Zhang et al., 2022), through which tasks such as semantic segmentation, estimation and other classification tasks are performed in order to perform tissue domain discovery(Lin et al., 2022; Ma et al., 2024; Yao et al., 2024). However, these methods often lack interpretability and testable parameters, presenting challenges in conducting multivariate spatial inference while providing clear descriptions of the data generating procedure. To bridge this gap, we developed integrated nested Laplace omics (INLAomics) (Fig. 1c), a comprehensive framework comprising: (i) a novel Bayesian hierarchical model utilizing a multivariate Gaussian Markov random field (GMRF) specification for (ii) efficient computational estimation to tackle modeling of spatial multi-omics data.

## 3 Results

### Development of INLAomics for the integration of multimodal spatial datasets

INLAomics infers protein abundance through a two-stage modeling strategy. First, gene co-expression programs are jointly modeled as general linear mixed models (GLMMs) using multivariate spatial autocorrelation Mardia (1988) (red plate Suppl. Fig. 1) with histological expert tissue annotations as covariates (green plate Suppl. Fig. 1). Second, protein abundance is modeled using the same covariates and plug-in estimates from the gene expression model, within a novel conditional spatial autocorrelation framework that extends the formulation of Jin et al. Jin et al. (2005) (red plate, Supplementary Fig. 1). Gene-to-protein effects are parameterized as both spot-specific and neighborhood-driven (i.e., neighbors-to-spot) contributions, defined per gene (blue plates, Supplementary Fig. 1). To enable higher-dimensional Bayesian inference, we then adopt a novel Multivariate Conditional Autoregressive (MCAR) formulation (Sec. 5) conducive to the INLA framework Rue et al. (2009) (Sec. 5 ). Specifically, INLAomics concurrently models gene and protein expression through a novel multivariate conditional CAR (MCCAR) structure (Sec. 5), where protein levels are conditionally dependent on multi-gene co-expression programs.. We provide INLAomics as an extension of the `R` software package `R-INLA` (Rue et al., 2017) which streamlines the implementation, usability and adaptation of our model and code, making them suitable for application to datasets and modalities beyond those considered here.

### INLAomics is computationally efficient in simulated settings

We first apply INLAomics in a simulated setting to show that our approximation provides similar results as to those obtained when no approximations were applied. We simulated data obtained from a real spatial multi-omics experiment (Vickovic et al., 2022) (murine spleen data, n=341 grid spots) and modeled parameters in order to compare between estimates recovered with INLAomics and those obtained with Markov Chain Monte Carlo (MCMC)(Sec. 5). Our sequential estimation procedure within the `R-INLA` environment (Rue et al., 2017) significantly reduced the estimation time (from days to minutes) as compared to MCMC methods implemented in `Stan` (Stan Development Team, 2023)(Sec. 5) while estimating the main parameters of interest that infer mRNA to protein abundance to a similar degree of accuracy (Suppl. Fig. 2a). We additionally show that traditional approaches, such as pairwise correlation analysis, fail to capture the complexity of mRNA–protein relationships, particularly in spatially resolved settings, underscoring the need for models that account for both spatial structure and multi-modal dependencies (Suppl. Fig. 2b).

### INLAomics identifies gene co-expression programs to build gene-to-protein networks

Secondly, we explored how INLAomics performs in real experimental settings. In a SPOTS(Ben-Chetrit et al., 2023) spatial multi-omics experiment, a total of 19 mRNAs (88.48*±*35.93 UMIs) and protein (1217.06*±*409.31 counts) pairs were spatially profiled by sequencing in a total of n=5,412 grid spots (Sec. 5, Suppl. Fig. 3). Previously(Ben-Chetrit et al., 2023), protein expression, as the modality detected at a greater abundance, was used as the driving factor in either cell type or tissue domain detection (Suppl. Fig. 3a), while the mRNA information was used *post hoc* to define cell-specific states within the clusters or domains (Suppl. Fig. 3b,c). Additionally, the relationship between mRNA and proteins was investigated at the level of pairwise correlations, where the correlation metric was mostly driven by count dropouts in either of the two modalities (Suppl. Fig. 3d, e).

To address these limitations, INLAomics fits a series of mRNA–protein models to iteratively construct a gene-to-protein network. The baseline model (*G* = 1) includes only a single mRNA–protein pair. Additional mRNAs are incorporated one at a time (*G >* 1), with model parameters re-estimated at each step. The inclusion order is determined by the absolute magnitude of the regression coefficients when gene expression levels are treated as covariates, prioritizing those with largest effect (Sec. 5). Finally, model selection is based on the deviance information criteria (Spiegelhalter et al., 2002) (DIC), adding genes until the DIC no longer decreases.

We highlight two illustrative cases comparing INLAomics with traditional correlation-based methods in estimating the significance of mRNA–protein relationships. In both approaches, F4/80 protein abundance was not significantly associated with its encoding gene, Adgre1 (Fig. 1d, left; Supplementary Fig. 3e). However, INLAomics identified a significant association between F4/80 protein levels and a distinct network of genes not coding the F4/80 protein—Cd163, MadCam1, and Siglec1—revealing putative co-regulatory or coexpression patterns resulting from cell-cell interactions otherwise missed by pairwise correlations. A second example involved the CD4 protein where standard correlation analysis showed a weak and non-significant association with Cd4 mRNA but INLAomics identified a coordinated expression program involving Cd4, Cd3e, Cd8a, and Cd163 that significantly influenced CD4 protein abundance (Fig. 1d, left; Supplementary Fig. 3e). These examples underscored the enhanced sensitivity of INLAomics in capturing biologically meaningful relationships beyond direct gene–protein pairs.

Interestingly, when incorporating spatial neighborhood effects, we observed that mRNA expression in neighboring tissue regions also influenced protein abundance at a given spatial location (Fig. 1d, right). For F4/80, Adgre1 and Siglec1 transcripts in neighboring spots significantly contributed to protein expression, despite Adgre1 showing no local effect in standard models. Similarly, for CD4, the spatial expression of Cd163, Cd3e, and Cd4 in the surrounding microenvironment was significantly associated with CD4 protein levels. These findings highlight the added value of spatially aware modeling in uncovering non-local mRNA co-expression patterns that influence protein expression.

Together, these results demonstrate that INLAomics robustly infers gene-to-protein networks, revealing that protein abundance is shaped by multi-gene expression programs and influenced by spatially localized mRNA dynamics. With this, INLAomics addresses key limitations of correlation-based methods and offers a more comprehensive framework for dissecting gene–protein relationships within tissue microenvironments.

### INLAomics for predicting tissue-level protein expression from gene co-expression programs

To evaluate the predictive performance of INLAomics, we trained the model under two conditions: (I) using protein measurements alone (*G* = 0), and (II) incorporating gene-protein co-expression programs (*G >* 0). For the latter, we selected the 200 most highly expressed genes as inputs, as they typically exhibited a greater degree of spatial variability and contributed meaningfully to the protein GMRF (see Methods, Sec. 5). Protein expression estimates for F4/80 and CD4 from gene–protein networks closely matched those from transcripts-per-million (TPM). Notably, spatially structured components captured by INLAomics explained substantial spot-level variance, with TPM concordance increasing as more genes were integrated into the networks (Suppl. Fig.4). To obtain protein predictions, the model parameters were then estimated using approximately 90% of the data, and predictive accuracy was assessed on the remaining 10% of spatial spots from the same tissue section, encompassing all four annotated tissue domains (Fig. 1e). Compared to the baseline model relying solely on protein data, prediction accuracy improved consistently when incorporating multiple genes (Fig. 1f, Suppl. Fig. 5, Suppl. Tab. 1). Furthermore, we demonstrated the generalizability of INLAomics by applying it in a whole-slide tissue prediction task. In this case, model parameters were trained on a single tissue section and subsequently applied to an independent section not included in training. Consistent with prior results, the model exhibited improved predictive accuracy for protein expression when leveraging multi-gene networks. (SPOTS murine spleen, Replicate 2, *n* = 2883 spots) (Suppl. Fig. 6). Together, these results demonstrate the capacity of INLAomics to accurately predict spatial protein expression from spatial transcriptomics data in tissue sections where antibody-antigen interactions had not been directly measured.

To test INLAomics in more challenging scenarios outside of the previously profiled mouse model system, we applied INLAomics to clinically relevant tissues: heathy human tonsil and breast cancer(Ben-Chetrit et al., 2023). When applied to human tonsil data (n = 4,908 grid spots), we again observed substantial improvements in protein prediction accuracy relative to the baseline model (Suppl. Fig. 7). This scenario underscored the significant implications of INLAomics, where gene expression alone can be leveraged to predict spatial protein profiles in tissue sections when expert tissue annotations are not available as inputs to the model. Next, we tested the predictive power of INLAomics in a setting of breast cancer data (n = 1,978 grid spots)(Ben-Chetrit et al., 2023). In cancer tissues, as compared to the spleen or tonsil examples above, limited tissue structure remains which impacts the overall spatial variability of proteins and genes present, and many of the genes exhibit low expression levels to meaningfully contribute to the conditioning set. In this case, when the tissue was unstructured, we randomly subset approximately 10% test spots from the tissue to repeat a similar testing procedure as described above. We observed that conditioning on more than one gene was generally not favored, and that it only marginally improved the model results as compared to the unconditional model (Suppl. Fig. 8, Suppl. Tab. 3). Conceptually, the spatial structure in the protein CAR model emerges from a combination of random and spatial noise (Sec. 5). Conditioning on gene expression can help account for spatial autocorrelation in the protein signal; however, its effectiveness hinges on the spatial heterogeneity and predictive power of the selected genes — which, in this case, were limited.

Overall, INLAomics performs effectively in structured tissues, where it leverages higher levels of gene expression and spatial patterns to predict protein profiles with high accuracy, even without expert annotations. In unstructured cancer tissues, however, the lack of spatial structure and low gene expression levels result in only marginal improvements over the baseline model. Nonetheless, in both cases, the interpretability and transparency of our method allow for a rapid selection of gene-to-protein networks in a specific context.

## 4 Discussion

A central challenge in multi-omics research lies in the weak concordance between RNA abundance and protein expression, driven by layers of post-transcriptional regulation. Existing computational methods—including deep learning, Gaussian processes, and dimensionality reduction—frequently overlook spatial dependencies, which are essential for accurately modeling tissue-resident cellular behavior. To address this, we present INLAomics, a Bayesian hierarchical framework for the integrative analysis of spatial transcriptomics and spatially resolved proteomics data.

INLAomics employs a MCCAR model and INLA estimation to identify gene-to-protein relationships that are often overlooked by correlation-based methods. The model explicitly incorporates spatial effects providing improved accuracy in estimating protein abundance and understanding gene-to-protein relationships in the context of tissues. By jointly modeling multiple transcripts, we demonstrate enhanced prediction of protein levels compared to approaches based on protein measurements alone or single-gene models. These results suggest that coordinated analysis of gene expression and protein biosynthesis uncovers putative co-expression and regulatory interactions that may be obscured when proteins are considered in isolation. Moreover, examining shifts in covariance structure upon the inclusion of multiple genes offers mechanistic insight into the drivers of protein expression.

While INLAomics offers substantial computational efficiency—scaling well with both sample size and model complexity—it is constrained to models with Gaussian latent fields. The method’s reliance on sparsity in the spatial covariance matrix, though beneficial for scalability, imposes limitations on capturing long-range or highly non-local spatial dependencies–a limitation that could be addressed in future work.

## 5 Methods

### Spatial multi-omics data

SPOTS fresh frozen murine spleen and murine breast cancer data were downloaded from the GEO using accession no. GSE198353. FFPE spatial proteogenomics data on human tonsil was downloaded with the 10X Genomics Datasets website Visium10x datasets. The slide for the simulation study was downloaded from Broad’s Single Cell Portal with accession no. SCP979. Processed count matrices were used as provided by the authors without any alterations.

### INLAomics model specification

We denote the vector of measurements for gene/protein *i* by 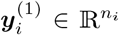 and the corresponding vector of RNA molecules by 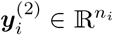 . We assume the measurements for gene/protein *i*, spot *j* and assay *ℓ* follows a Poisson distribution

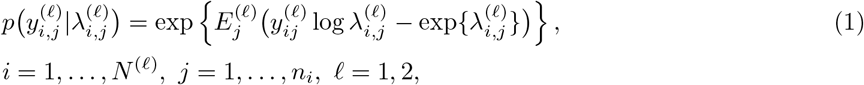

where 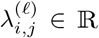 is the rate parameter. 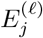 is a spot specific size factor treated as known and set to (Maniatis et al., 2019)

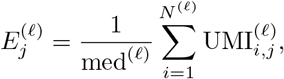

where 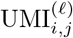 denotes the number of unique molecular modifiers for gene/protein *i*, spot *j* and assay *ℓ*, and med^(*ℓ*)^ is defined to be the median sequencing depth of the UMIs of assay *ℓ*. It is assumed that 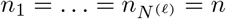 for *ℓ* = 1, 2.

The rate vector 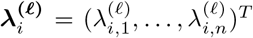 of (1) is modelled as a spatial generalized linear mixed model (GLMM),

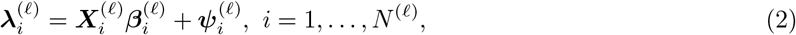

where 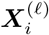 is a *n* × *p* model matrix and 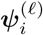 is a GMRF. The GMRF covariance is modeled as a mixture between independent noise and an intrinsic CAR process (Leroux et al., 2000)

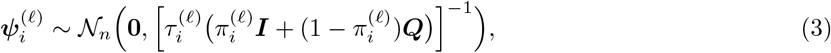

where 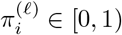 and ***Q*** is the *n* × *n* matrix with elements

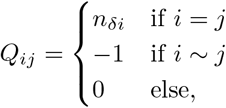

where *n* _*δi*_ is the number of neighbors of spot *i* and *i* ∼*j* implies that spots *i* and *j* are neighbors in the graph of the GMRF. We shall denote 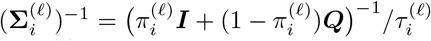 in the sequel.

The dependence of processes ***y***^(1)^ and ***y***^(2)^ will be introduced on the level of the GMRFs as a conditional distribution 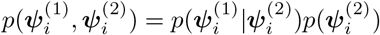 defined as (Jin et al., 2005)

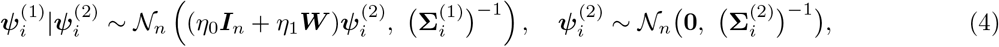

where ***W*** is a neighbourhood matrix of the spots, ***I***_*n*_ is the *n* × *n* identity matrix, *η*_0_ ∈ ℝ measures the spot-to-spot effect of RNA on that of protein and *η*_1_ ∈ ℝ measures the effect of the neighboring RNA measurements on the protein.

As RNA are generally measured over a much larger set of genes in comparison to number of proteins, i.e., *N* ^(1)^ ≪ *N* ^(2)^, it is of interest to consider forms of MCAR error terms where multiple RNA genes affect a single protein. As such, consider the following extension of (4) to include that of *G* ∈ {1, … , *N* ^(2)^} genes,

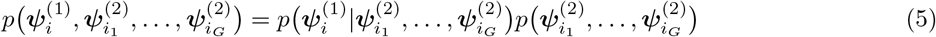

where *{i*_1_, … , *i*_*G*_*}* is a subsequence of the genes and

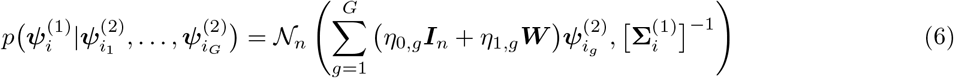

and

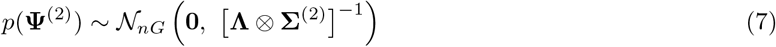

where 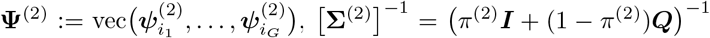 and **Λ** is a R × R positive definite matrix defined as

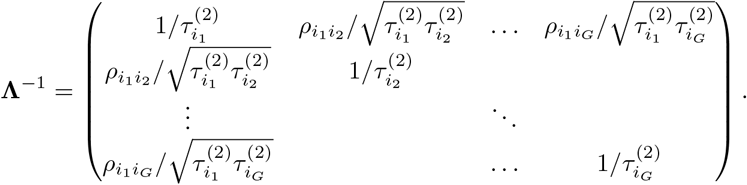

The Kronecker structure of (7) follow the specification of Mardia (1988), where we have assumed that 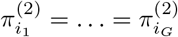. The conditional specification of (6) and (7) implies the joint distribution

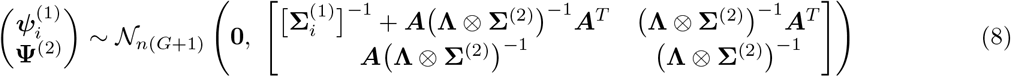

where ***A*** is a *n* × *nG* block-matrix defined as

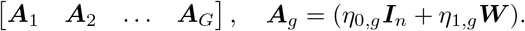

Note that if we model proteins and RNA jointly as (7) with, e.g., *G* = 2, we have that it arises as a special case of (8) assuming *π*^(1)^ = *π*^(2)^ and (*η*_1,1_, *η*_1,2_)^*T*^ = **0**. They are then functionally related as 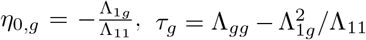, for *g* = 1, 2, *τ*_1_ = Λ_11_ and 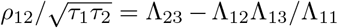. Note that Λ_*ij*_ is the element (*i, j*) of **Λ** implied by (7), for the joint of one protein in the first coordinate and *G* = 2. A similar result for *G* = 1, and *ρ*_12_ = 0, was noted in Jin et al. (2005), and follow from a standard result in matrix theory, see Corollary 8.15.12 of Harville (1997).

For both *G* = 1 and general *G*, Bayesian estimation procedures are considered so priors need to be specified to complete the model. For general *G*, which shall be referred to as the many-to-one case, we consider the following priors for the scale parameters

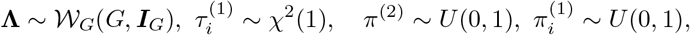

where 𝒲_*p*_(*r*, **Σ**) denotes the Wishart distribution where **Σ** is a *p* × *p* positive definite matrix and *r > p*− 1. In the case where the off-diagonals of **Λ** are 0, the Wishart specification simplfies to independent chi-square distributions with 1 degree of freedom, denoted by *χ*^2^(1). For the one-to-one case, *G* = 1, the only difference is for **Λ** where we instead have 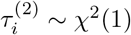. For the location parameters, we set

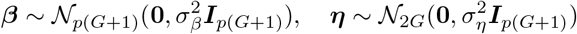

where 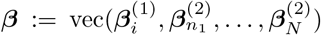 and ***η*** = (***η***_0_ , ***η***_1_ )^*T*^ . Unless stated otherwise, we set 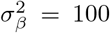 and 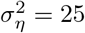.

Let ***θ***_1_ and ***θ***_*G*_ denote the sets of hyperparameters of the spatial error terms in the one-to-one and many-to-one case respectively. It is then assumed that the observations are independent conditional on the spatial error term, so full posterior for the many-to-many case is given by

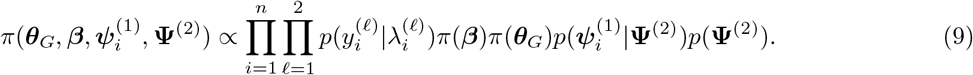

Similarly, for the one-to-one case

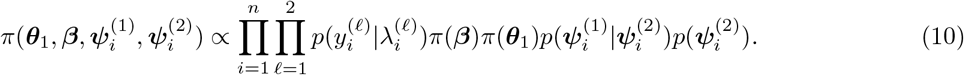

For estimation of both (9) (10), we shall consider integrated nested Laplace approximation (INLA) of Rue et al. (2009) utilizing the R(Stan Development Team, 2023) package R-INLA. Such approximations have greater computationally efficiency relative to Markov chain Monte Carlo (MCMC) methods, making them particularly useful in the spatial multimodal setting. The implementation of proposed MCCAR is implemented in `R-INLA` via the `inla.rgeneric` function which allows for the implementation of new GMRF models.

We give a brief outline of the estimation procedure in the one-to-one case, and refer to Rue et al. (2009) and Rue et al. (2017) for further details. First, denote the joint prior distribution of ***β***^(*ℓ*)^ and ***ψ***^(*ℓ*)^ by ***z***^(*ℓ*)^, which will be normally distributed

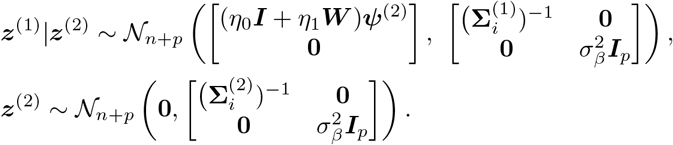

Subsequently, partition ***θ***_1_ as 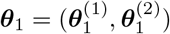, where 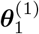 denotes the hyperparameters of the RNA processes and 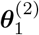 those of the protein process. We then approximate the posterior of ***θ***_1_ as

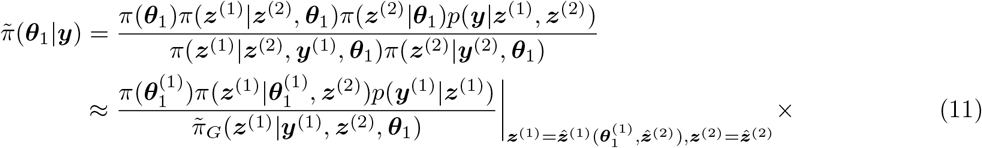

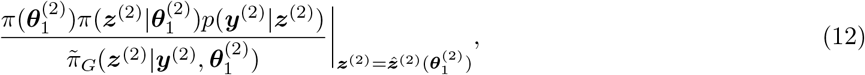

where 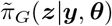 denotes the Gaussian approximation of the conditional distribution of ***z***, and 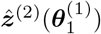 denotes the mode of the conditional distribution of ***z***^(2)^ for a given 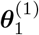 and 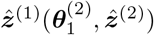 is defined equivalently. The protein process (11) can be seen as an empirical Bayes procedure, where the dependency of ***z***^(1)^ on ***z***^(2)^ in the denominator of (11) is disregarded to obtain the estimate 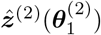. This approach is necessary to enable the use of `R-INLA` for the conditional GMRF, as (8) is not applicable in `R-INLA`, and implies that all parameters of the RNA model can be estimated first and then plugged into (11). We note that a thematically similar plug-in estimation was utilized by Krock et al. (2021, 2023).

Assessing convergence of (12) and 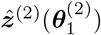 is crucial to understand the implications of the decomposition into (11) and (12). However, such efforts are challenging as the dimension of the GMRF increase at the same rate as the sample size. For a detailed discussion, we refer to Rue et al. (2009). Another issue with the decomposition is the compounding of error rates of the Laplace approximations. If the true latent fields ***z***^(1)^ and ***z***^(2)^ converge to degenerate normal distributions of rank *q*_1_ and *q*_2_, respectively, as *n* → ∞, the error rate becomes of the order 𝒪 (*q*_1_*/n*) 𝒪 (*q*_2_*/n*) = 𝒪 (*q*_1_*q*_2_*/n*^2^). Similarly, if the components of the latent fields are independent, the error rate is unaffected by the decomposition and the error rate is 𝒪 (1). To address these issues, a small simulation study is conducted as a sanity check, comparing the INLA-based sequential procedure to MCMC implemented in Stan(Stan Development Team, 2023).

Lastly, the extension to the many-to-one case only concerns (12), where we will utilize a modification of the the `R` package `INLAMSM` (Palmí-Perales et al., 2021), which implements (7) using a different covariance structure. All analysis was carried out in R (R Core Team, 2021) V. 4.3.1.

### Comparison to MCMC

We consider a Monte Carlo (MC) simulation study, based on 10^4^ replications, where the suggested procedure is compared to MCMC implemented in Stan (Stan Development Team, 2023). Focus is on estimation of the hyper-parameters of the conditional GMRF in (11), rather than biologically relevant tissue. We consider spots on even a grid of 341 spots. The GMRF are generated as (4),

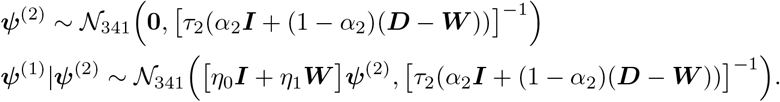

where we set *τ*_1_ = *τ*_2_ = 1, *η*_0_ = 0.5, *η*_1_ = −0.25 and *α*_1_ = *α*_2_ = 0.2 Subsequently, the data is generated as

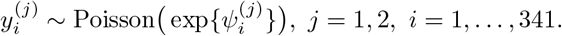

Posterior samples are generated via Stan Development Team (2023) with a total of 2 *·* 10^4^ samples generated with the first 10^4^ discarded as warm-up. The simulations where carried out in R, where the average estimation time of MCMC is 20 minutes and that of the proposed procedure is 12 seconds. The span of computation time for MCMC is large, taking anywhere from 13 minutes up to 40 minutes. Note that in the MCMC procedure, the log determinant of the GMRF is computed efficiently as

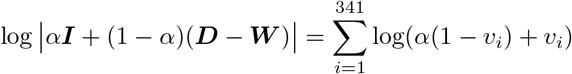

where ***v*** are the eigenvalues of ***D*** − ***W*** . Thus the time difference is not due to inefficient implementation of the MCMC procedure. For estimation `R-INLA` V. 23.12.17 was used and for MCMC, `Stan` V. 2.26.1 with `R` interface `R-stan` V. 2.26.23.

### Comparison to univariate correlation

We consider a MC simulation study, based on 10^4^ replications, illustrating the potential pitfalls of univariate measurements in the presence of complex multivariate correlation structures. The same grid of 341 spots utilized in preceding section is utilized and data is generated as follows

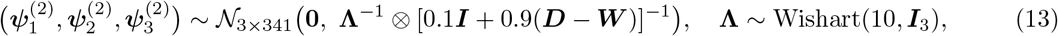

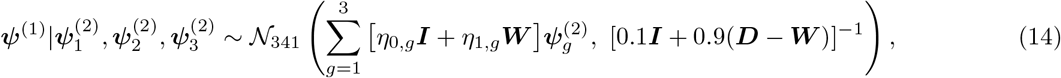

where the cross-assay parameters are defined as

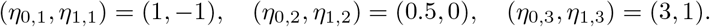

We utilize correlation as univariate measurement of associations between (13) and (14). We note, in particular, an issue with the correlation between 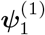 and ***ψ***^(2)^, which is estimated to be close to 0, i.e. averaging between the effects *η*_0,1_ = −*η*_1,1_ (Suppl. Fig. 2b).

### Model selection

For estimation, we start with the RNA model where the joint log rates of *g* = 1, … , *G* genes are modelled as

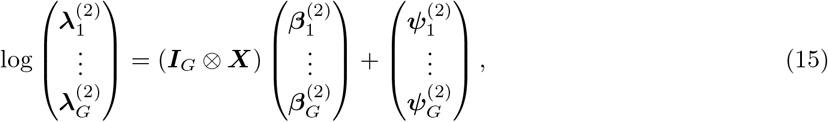

where ***X*** is a *n* × *p* matrix encoding the cell annotations and 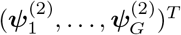 is distributed as (7). Note that (15) simplifies to

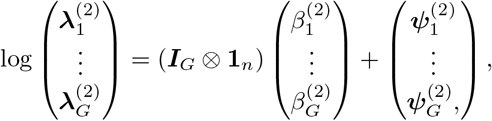

when no cell annotations are used, where **1**_*n*_ is a *n* × 1 vector of ones.

For each protein, we start by fitting the conditional GMRF (4) using only the protein and gene pair, i.e., *G* = 1. Subsequently we add one gene at the time and re-estimate the model. The order at which genes are added is based on the absolute value of the corresponding fixed effects coefficients when the gene expressions are included in the model matrix, with an unconditional CAR error structure. Then, the *G* = 1 and *G* = 2 models are compared using the DIC, defined as

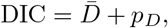

where 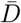 is the posterior expected deviance and *p*_*D*_ is the “effective” number of parameters. For DIC, lower values are favourable, so if the DIC decreases for the expanded model, the procedure is repeated of model comparison between *G* = 2 and *G* = 3. The process is repeated until DIC increases, and the preceding model is selected.

### Analysis of gene-protein programs in the mouse spleen

For the analysis of gene-protein programs, we utilized two replicates of murine spleen spatial multi-omics data (Ben-Chetrit et al., 2023). The dataset contained a total of 21 gene and protein pairs however two pairs were dropped from the analysis (Ly6C and Ly6G) as genes are not sufficiently expressed to provide meaningfully. The authors additionally state in their work that the Ly6C protein did not work. Means refer to estimated posterior means and significance (solid dots) is interpreted as the 95% credible set not covering 0, the necessary estimates are obtained directly from `summary.hyperpar` in the return object of `inla()`.

For correlations, we first normalized the data using `Seurat` V. 4.4.0, `NormalizeData()` and `ScaleData()`. These normalized counts were used to generate plots in Suppl. Fig. 3 where, further, we report Pearsons’s r correlation via `cor()` and the absolute value of the coefficients from regressing genes on proteins with a CAR error from, estimated via `inla()`. These can obtained `summary.fixed` from the return object of `inla()`.

### Protein prediction analysis

For predictions (Fig. 1 and Suppl. Fig. 4, 8, 7), TPMs are computed by normalizing the raw counts to sum to 10^6^ across genes and proteins separately. Further, ADTs are scaled across spots to have mean 0 and variance 1 using `scale()`. Note further that there is no `predict()` function for `R-INLA`, as predictions are done simultaneously as model building. Test data is predicted by setting the corresponding measurements to NA, and are obtained through the `summary.fitted` element of the `inla()` return object.

Constrained parameters has to be transformed to the real line to apply Laplace approximations. To obtain the parameters *π* and *σ*^2^ in Suppl. Tab. 3, we use `inla.tmarginal()` to convert estimates to the original scale. It takes the inverse mapping of the aforementioned transformation as argument, which are *e*^−*τ*^ for the variance and 1*/*(1 + *e*^−*π*^) for the convolution parameter.

## Code availability

All code has been deposited and made available on GitHub (https://github.com/nygctech/INLAomics).

## Acknowledgements

The computations were enabled by resources provided by the National Academic Infrastructure for Supercomputing in Sweden (NAISS), partially funded by the Swedish Research Council through grant agreement no. 2022-06725. Work was supported by the Knut and Alice Wallenberg Foundation, Beijer Laboratory for Gene and Neuro Research, Science for Life Laboratory, Target ALS, Bill and Melida Gates Foundation, the MacMillan Center for the Non-coding Genome, 1U54 AG076040-01 and 1RM1 HG011014-01 (S.V.). S.V was supported as a Wallenberg Academy Fellow and SciLifeLab Fellow at Uppsala University.

## 6 Author contributions

L. A. and S. V. conceived and designed the study. L. A. developed the model with guidance from S. V. L. A. and S. V. interpreted the data, discussed the results and wrote the manuscript.

## 7 Competing interests

The authors declare no competing interests.

---

**Suppl. Fig. 1:**
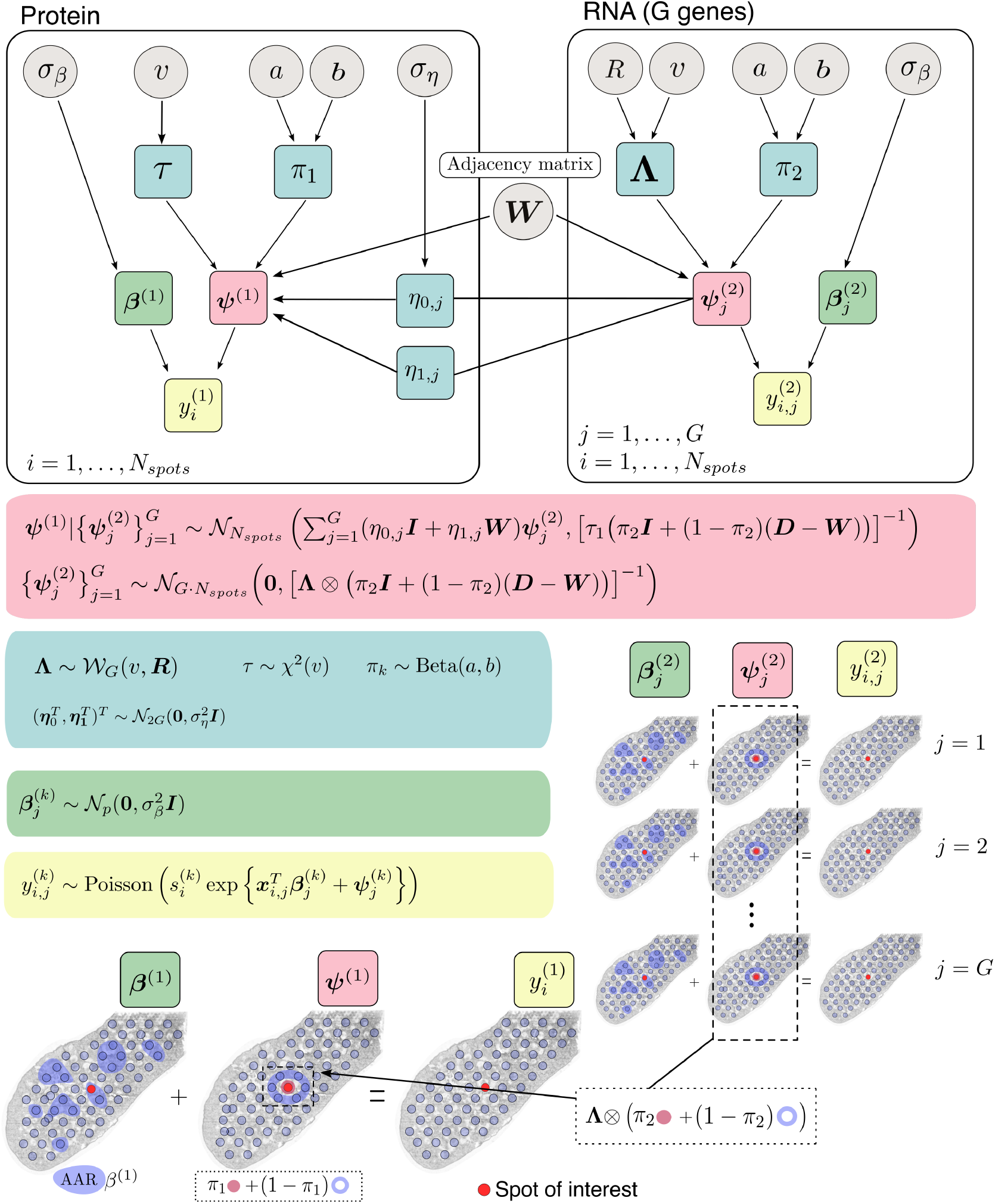
Graphical representation of the INLAomics model using plate notation. Grey circles denote hyperparameters with fixed values, while random components are listed within squares. Colored regions highlight distinct model components: spatial effects (red), prior distributions of spatial effects components (blue), measurement models (yellow), and fixed effects (green). Directed edges indicate the hierarchical structure of the model. All model distributions are explicitly specified. A biological schematic accompanies the plate diagram, using matched colors for interpretability. INLAomics jointly models expression across *G* genes with a Kronecker-structured dependency between the genes (second row, red box). Estimated gene-level random effects 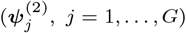 are propagated into the protein-level process (first row, red box). Spatial effects are modeled via convolutions between random fields and spatial noise structures, shown in dotted boxes.

**Suppl. Fig. 2:**
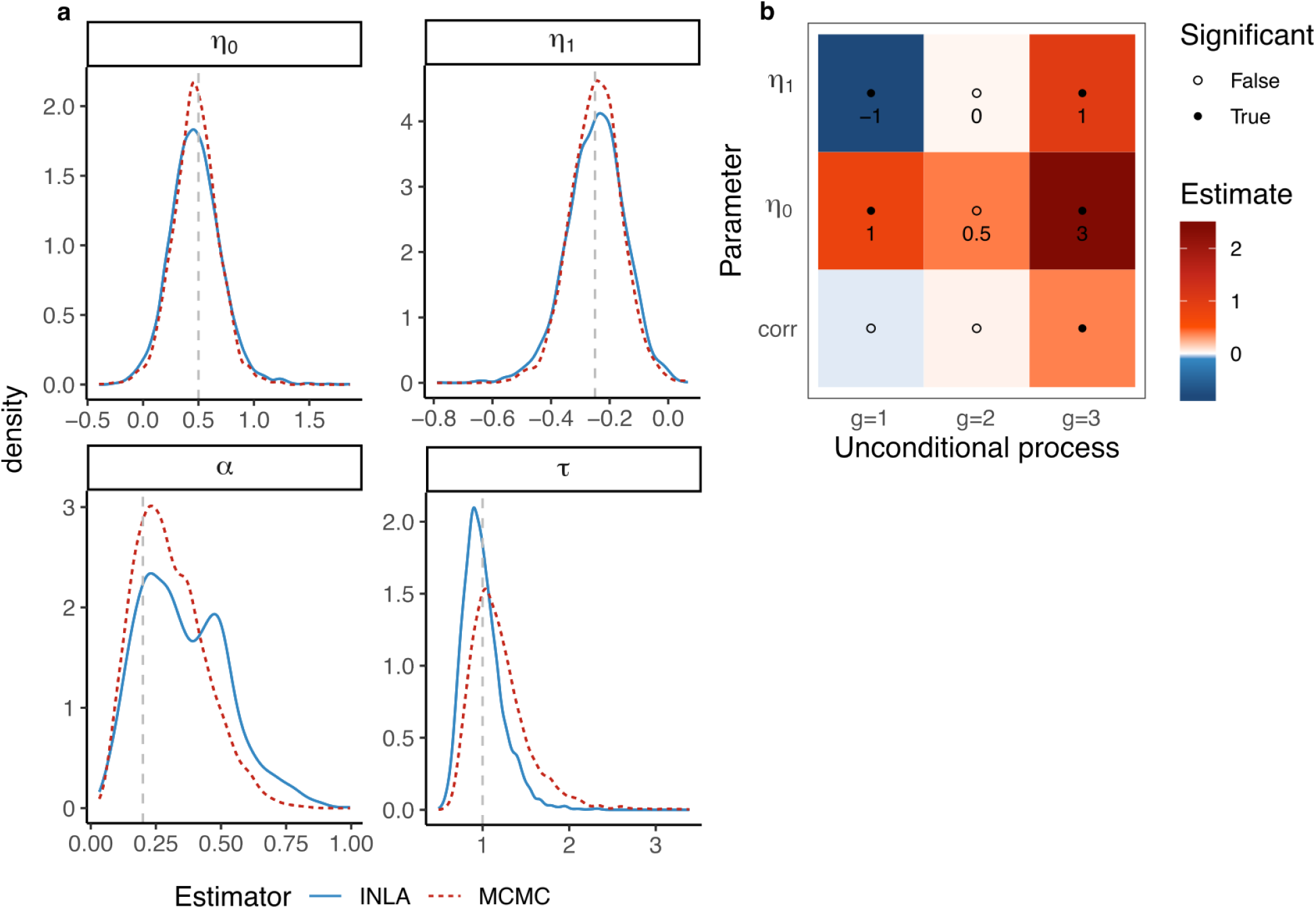
INLAomics in a simulated setting. **a**, Comparison of parameter estimates between MCMC (red dashed) and INLAomics (solid blue). *η*_0_ and *η*_1_ represent the spot-to-spot and neighbors-to-spot effects between assays. Additionally, for the protein process, *π* denotes the convolution between independent and spatial noise (*π* = 1 indicating no spatial structure) and *τ* denotes the precision (inverse variance). Grey lines lines mark the parameter values of the data generation process (Sec. 5 for details). **b**, Comparison of INLAomics and univariate correlations in measuring RNA-to-protein effects in simulated data (Sec. 5 for details). True values of cross-assay effects ***η***_0_ and ***η***_1_ are noted beneath the dots of rows 1 and 2 and estimated values are colored (color scale). Columns correspond to the indices of the unconditional process; for example, column 1 in rows 1 and 2 shows the cross-assay parameter estimates between simulated gene process *g* = 1 and protein, while row 3 displays the correlation (*corr*) between the processes. Columns *g* = 2 and *g* = 3 show equivalent estimates between protein and genes 2 and 3. Estimates significantly different from 0 are marked by solid dots.

**Suppl. Fig. 3:**
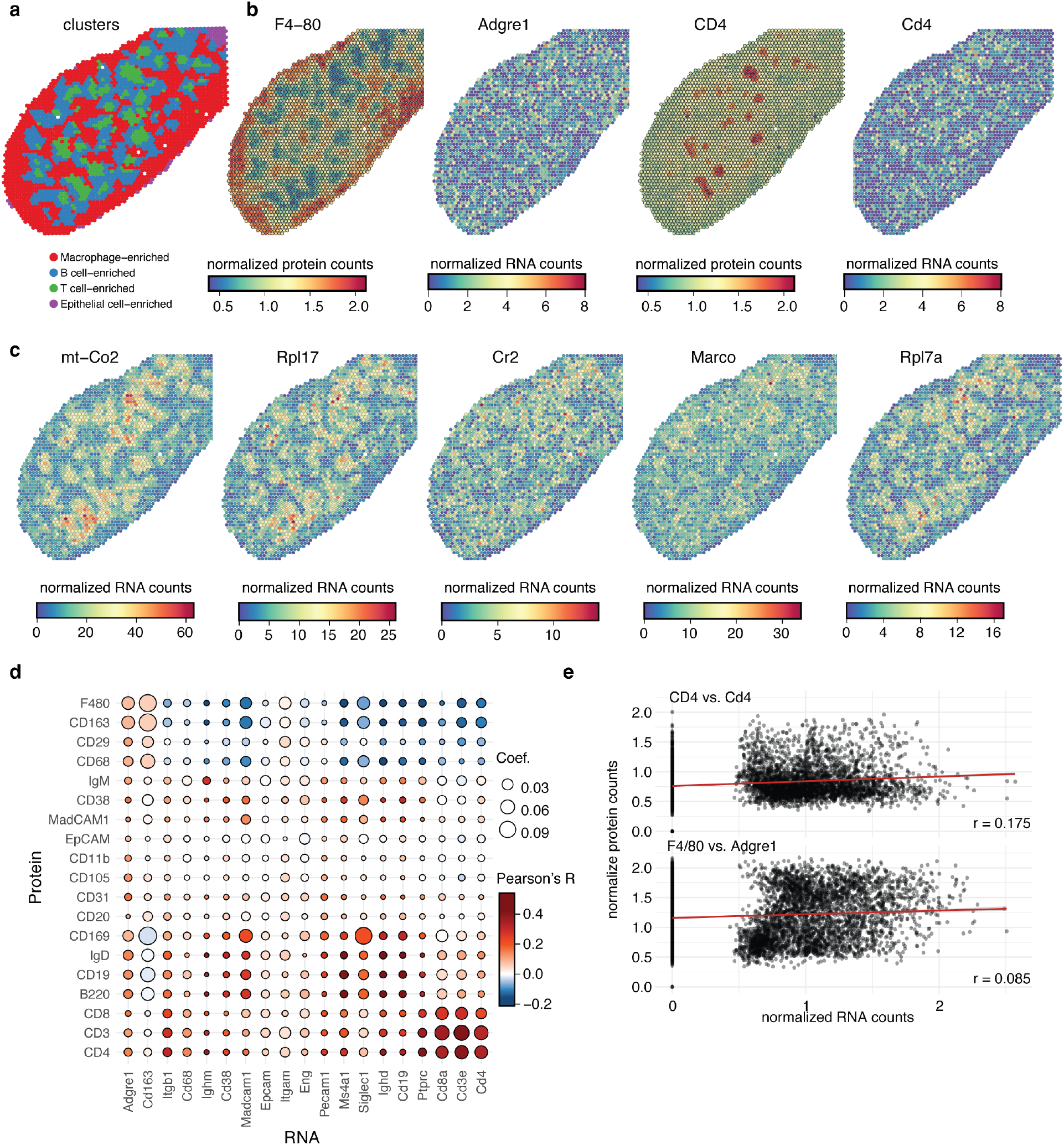
mRNA and protein counts analysis using a disjoint correlation model. **a**, Spatial clusters (color code) detected using protein-only counts data. **b**, Normalized expression (color scale) for two gene-protein pairs: F4/80 vs. Adgre1 and CD4 vs. Cd4. **c**, Normalized expression of genes detected as differentially expressed(Ben-Chetrit et al., 2023) in the B-cell enriched cluster in **(a). d**, Correlation plot (color scale) between normalized gene (columns) and protein (rows) counts with the linear correlation coefficient denoted for each pair (dot). **e**, Correlation plots for the two gene-protein pairs: F4/80 (y-axis) vs. Adgre1 (x-axis) and CD4 (y-axis) vs. Cd4 (x-axis). Pearson’s correlation coefficient (r) denoted in the right left corner for each respective pair. Red line represents linear regression.

**Suppl. Fig. 4:**
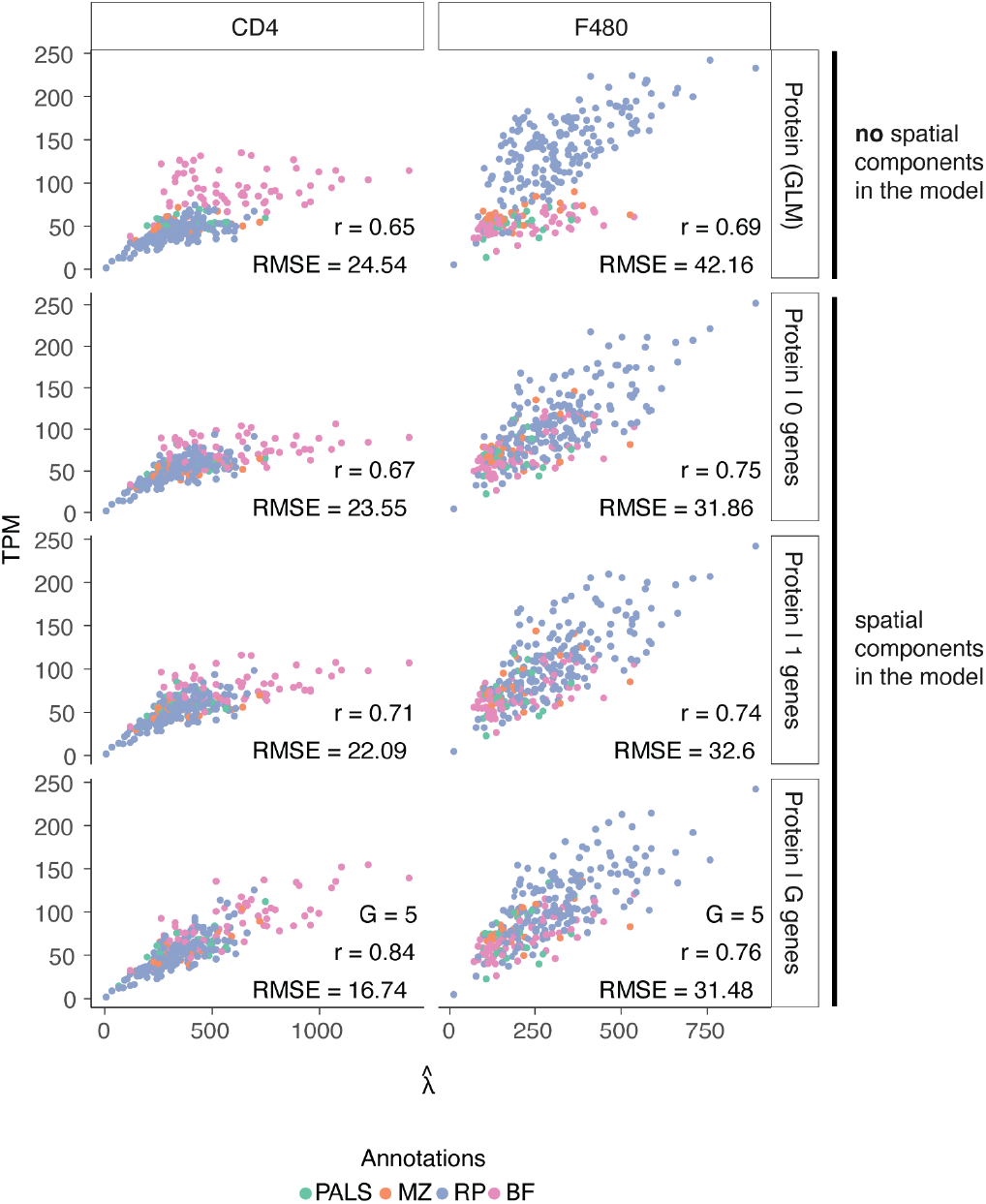
Inset prediction analysis for F4/80 and CD4 protein expression. Correlation (Pearson’s *r*) between TPM (*y*-axis) and estimated protein expression rates (*x*-axis) for all individual spatial spots (color code) used in the analysis. Color code denotes pathologist-informed tissue annotation categories. RMSE denoted for unconditional and multi-gene models (rows) for F4/80 and CD4 (columns). *G* denotes number of genes used in the model estimation.

**Suppl. Fig. 5:**
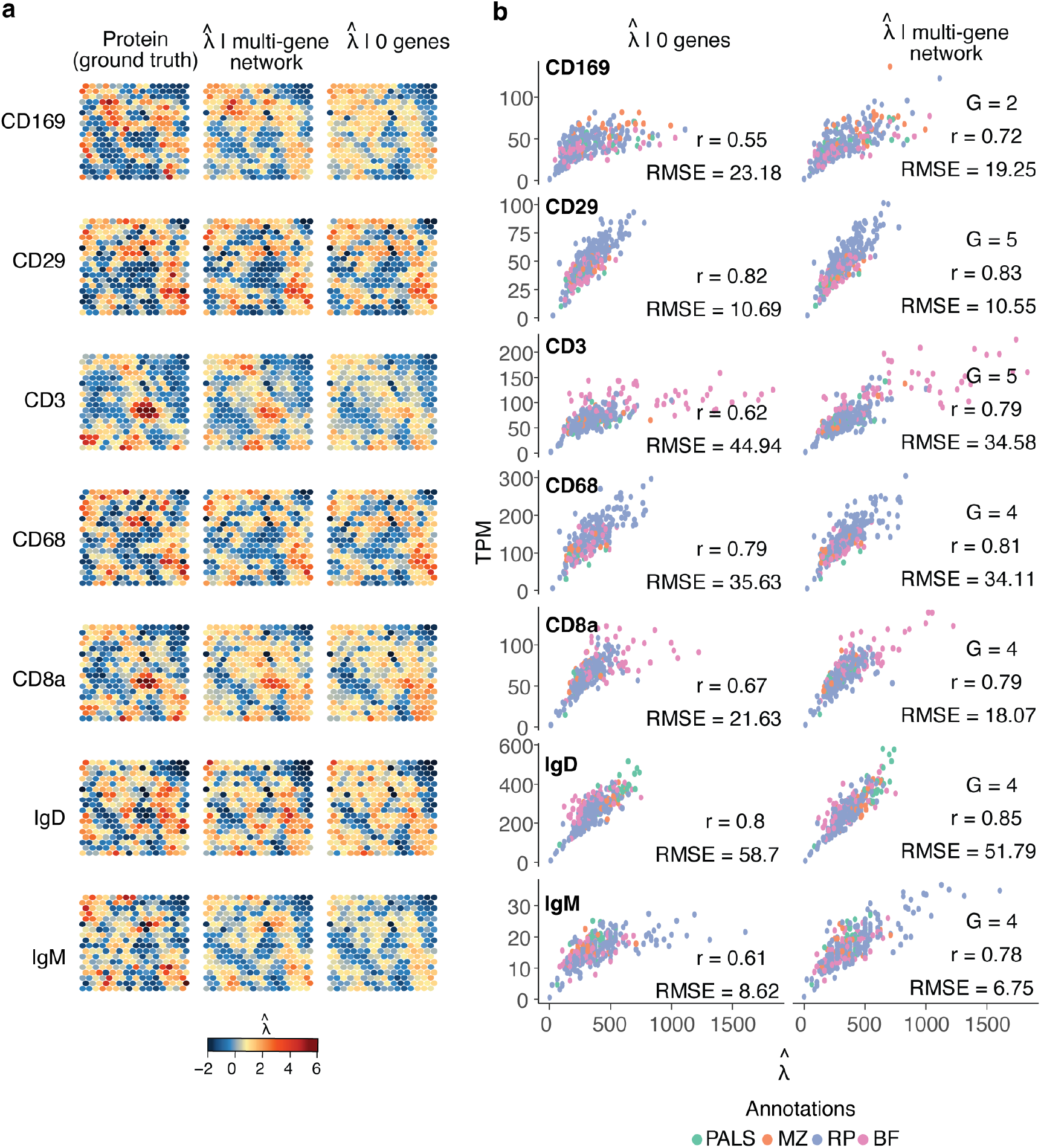
Inset prediction analysis for several gene-protein pairs. **a**, Comparison of predicted rates (color scale) for a selected subset of proteins (rows) between unconditional and conditional models (columns). **b**, Correlation (Pearson’s *r*) between TPM (*y*-axis) and estimated protein expression rates (*x*-axis) for all individual spatial spots (color code) used in the analysis. Color code denotes pathologist-informed tissue annotation categories. RMSE denoted for unconditional and multi-gene models (columns) for proteins (rows). *G* denotes number of genes used in the model estimation.

**Suppl. Fig. 6:**
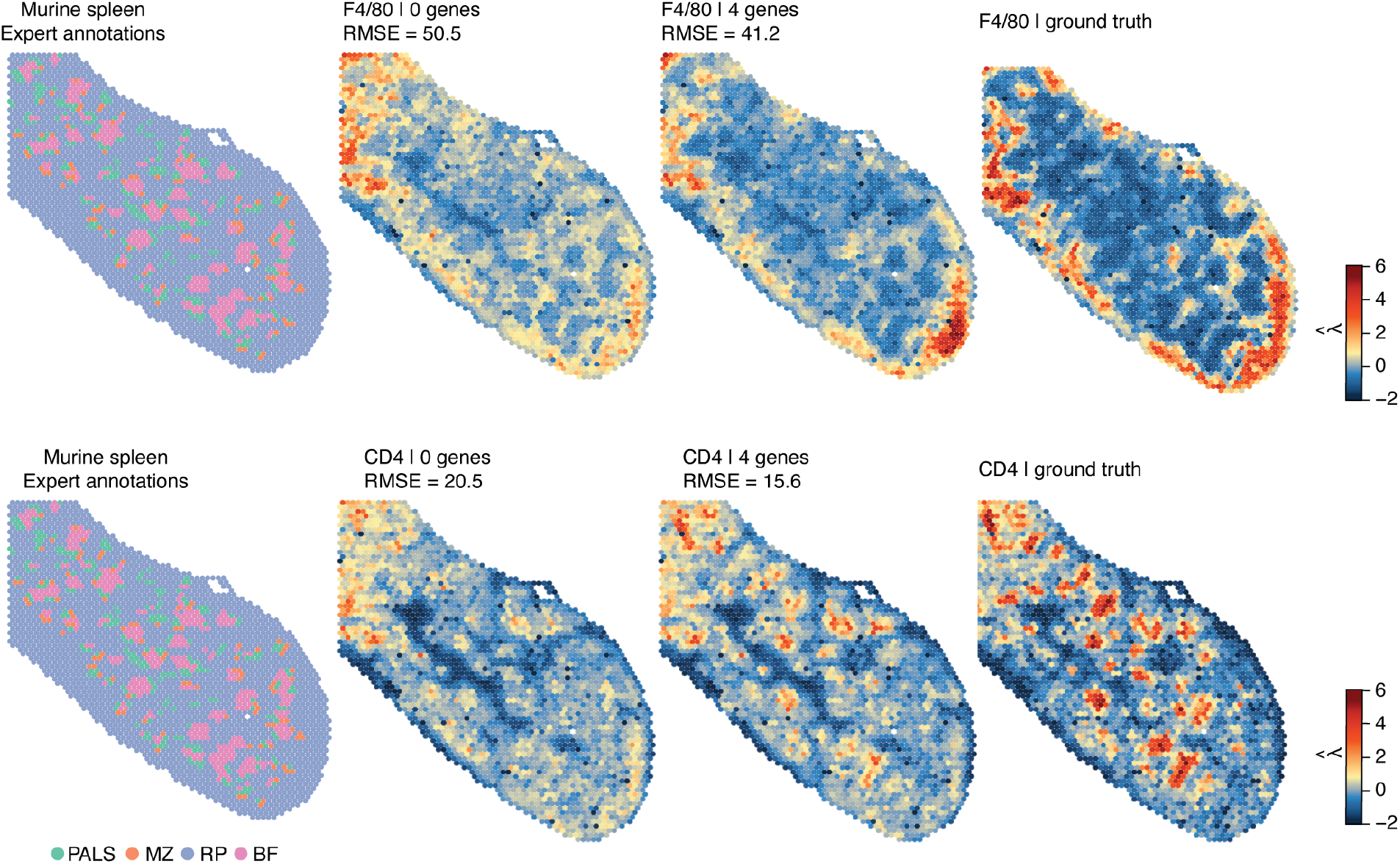
Whole slide prediction analysis for F4/80 and CD4 protein expression. Comparison of predicted protein expression rates (color scale) for whole slide spatial analysis of F4/80 and CD4 (rows) between unconditional and conditional models (columns). Pathologist annotations (color code) denoted on the far left.

**Suppl. Fig. 7:**
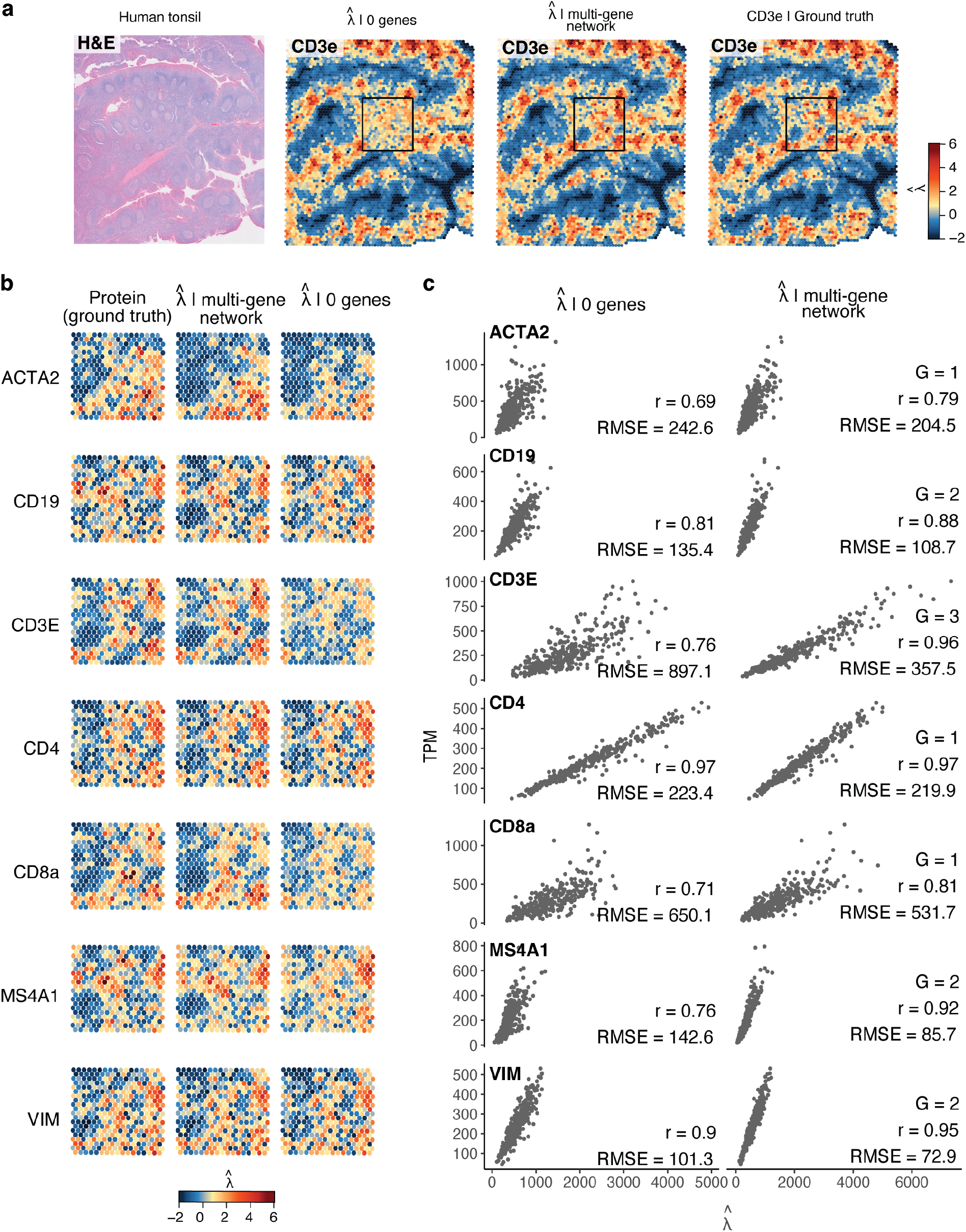
Inset prediction analysis for human tonsil. **a-b**, Comparison of predicted rates (color scale) for a selected subset of proteins (rows) between unconditional and conditional models (columns). **a**, Hematoxylin and Eosin (HE) stained tissue image of the human tonsil profiled with the spatial multi-omics technology and showing protein expression of CD3E (color scale) for the conditional, unconditional model and ground truth (columns). Test inset denoted (black rectangle). **b**, Predicted protein (rows) expression rates (color scale) for test inset marked in **a. c**, Correlation (Pearson’s *r*) between TPM (*y*-axis) and estimated protein expression rates (*x*-axis) for all individual spatial spots used in the analysis. RMSE denoted for unconditional and multi-gene models (columns) for proteins (rows). *G* denotes number of genes used in the model estimation.

**Suppl. Fig. 8:**
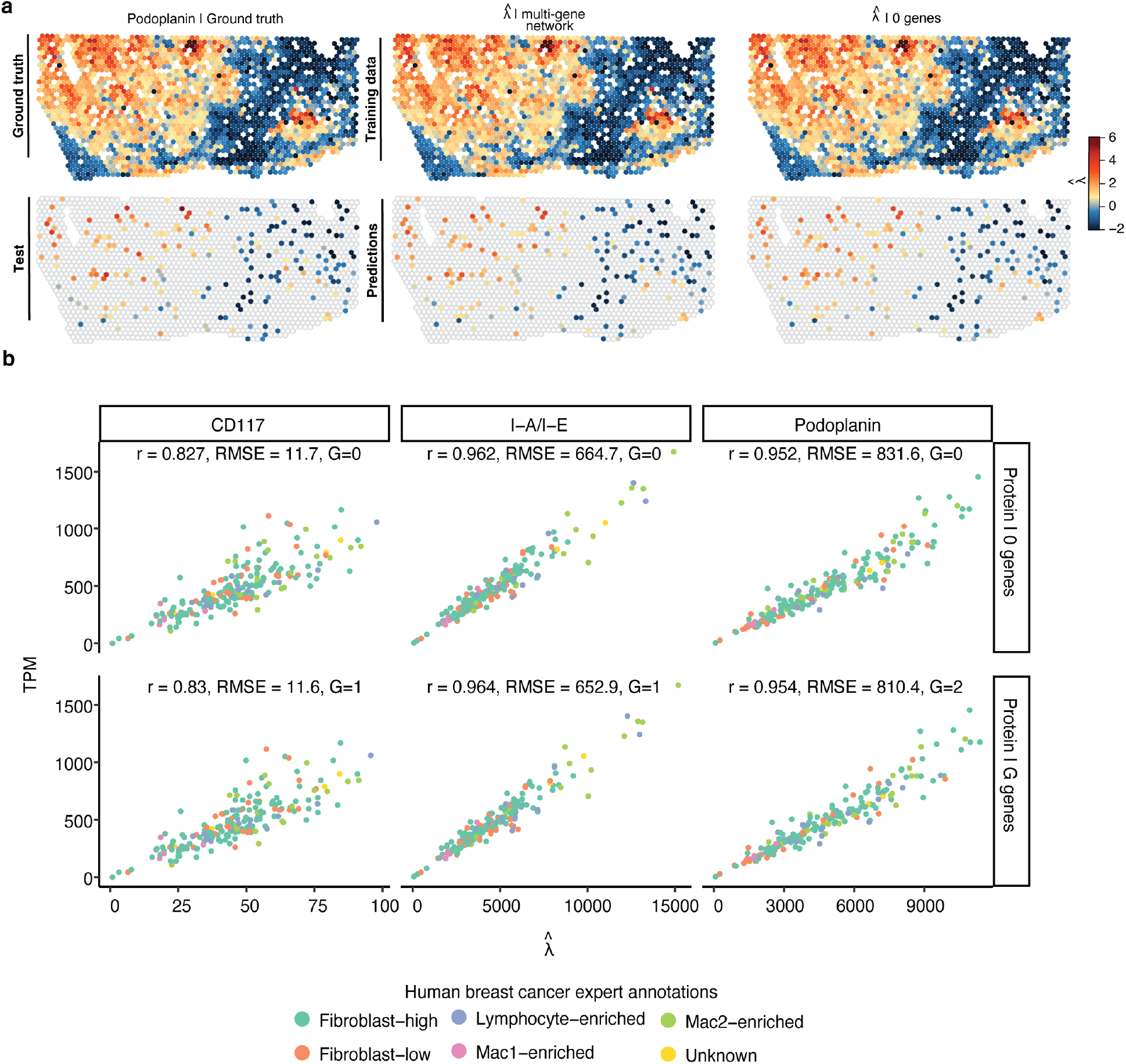
Prediction over randomly selected spots in spatial multi-omics breast cancer data. **a**, Comparison of predicted Podoplanin protein expression rates (color scale) to ground truth expression between unconditional and conditional models (columns). Prediction was carried out over a random subset of spatial spots. **b**, Correlation (Pearson’s *r*) between TPM (*y*-axis) and estimated protein expression rates (*x*-axis) for all individual spatial spots (color code) used in the analysis. Color code denotes pathologist-informed tissue annotation categories. RMSE denoted for unconditional and multi-gene models (rows) for CD117, I-A/I-E and Podoplanin (columns). *G* denotes number of genes used in the model estimation.

**Suppl. Tab. S1:** RMSE for subset of proteins (tonsil) with number of genes (G).

| Protein | RMSE |  |
| --- | --- | --- |
| B220 | 16.37 ($G = 0$) | 15.91 ($G = 3$) |
| CD105 | 8.02 ($G = 0$) | 7.86 ($G = 5$) |
| CD11b | 7.37 ($G = 0$) | 7.31 ($G = 3$) |
| CD163 | 16.37 ($G = 0$) | 15.84 ($G = 4$) |
| CD19 | 18.16 ($G = 0$) | 16.57 ($G = 4$) |
| CD20 | 3.83 ($G = 0$) | 3.77 ($G = 3$) |
| CD31 | 8.70 ($G = 0$) | 8.51 ($G = 4$) |
| Cd38 | 23.40 ($G = 0$) | 21.12 ($G = 4$) |
| EpCAM | 4.77 ($G = 0$) | 4.60 ($G = 2$) |
| MadCAM1 | 4.10($G = 0$) | 4.04($G = 4$) |

**Suppl. Tab. S2:** Model selection criteria (DIC) for gene-to-protein networks (spleen).

| Protein | Genes |  |  |  |  |  |  |
| --- | --- | --- | --- | --- | --- | --- | --- |
| F480 | Adgre1<br>43171.68 | Cd163<br>43168.32 | Siglec1<br>43161.74 | Madcam1<br>43160.76 | Ighd<br>NA |  |  |
| CD163 | Cd163<br>36613.94 | Madcam1<br>36601.28 | Eng<br>36599.88 | Siglec1<br>36586.46 | Pecam1<br>NA |  |  |
| CD29 | Itgb1<br>38132.84 | Cd163<br>38132.14 | Siglec1<br>38129.83 | Madcam1<br>38126.31 | Epcam<br>NA |  |  |
| CD68 | Cd68<br>45221.45 | Cd163<br>45215.72 | Madcam1<br>45212.73 | Itgam<br>NA |  |  |  |
| IgM | Ighm<br>31023.20 | Epcam<br>31018.66 | Cd3e<br>31011.79 | Cd68<br>31008.05 | Cd4<br>NA |  |  |
| CD38 | Cd38<br>44380.31 | Cd163<br>44378.32 | Siglec1<br>44359.95 | Cd8a<br>44334.80 | Eng<br>NA |  |  |
| MadCAM1 | Madcam1<br>28134.18 | Cd163<br>28137.75 |  |  |  |  |  |
| EpCAM | Epcam<br>26795.09 | Eng<br>26792.82 | Siglec1<br>26790.93 | Itgam<br>26792.59 |  |  |  |
| CD11b | Itgam<br>34852.51 | Epcam<br>34853.52 |  |  |  |  |  |
| CD105 | Eng<br>36228.06 | Cd4<br>36229.84 |  |  |  |  |  |
| CD31 | Pecam1<br>36796.43 | Eng<br>36795.00 | Cd8a<br>NA |  |  |  |  |
| CD20 | Ms4a1<br>28724.94 | Cd163<br>28724.28 | Itgam<br>28723.19 | Epcam<br>28728.76 |  |  |  |
| CD169 | Siglec1<br>38560.22 | Cd163<br>38554.61 | Cd8a<br>38544.91 | Madcam1<br>38540.48 | Cd3e<br>NA |  |  |
| IgD | Ighd<br>48782.75 | Itgam<br>48782.84 |  |  |  |  |  |
| CD19 | Cd19<br>39902.71 | Cd163<br>39898.36 | Cd8a<br>39872.84 | Madcam1<br>39866.78 | Ms4a1<br>39849.07 | Itgb1<br>39817.26 | Siglec1<br>NA |
| B220 | Ptprc<br>40103.21 | Cd163<br>40101.13 | Cd81<br>40068.49 | Eng<br>40068.09 | Madcam1<br>40043.31 | Siglec1<br>NA |  |
| CD8 | Cd8a<br>39850.64 | Cd3e<br>39825.94 | Cd4<br>39819.33 | Epcam<br>39805.61 | Madcam1<br>39806.33 |  |  |
| CD3 | Cd8a<br>40164.15 | Cd4<br>40125.44 | Itgb1<br>40113.02 | Madcam1<br>40108.97 | Cd163<br>40098.45 | Ptprc<br>NA |  |
| CD4 | Cd4<br>38669.83 | Cd163<br>38667.38 | Cd3e<br>38641.78 | Cd8a<br>38634.12 | Itgb1<br>38612.74 | Siglec1<br>NA |  |

**Suppl. Tab. S3:** Parameter estimates of selected models for the breast cancer data. Standard errors in parenthesis, *σ*^2^ = 1*/τ* represents the variance. *η*_0,*i*_ is the spot-to-spot effect of gene *i* and *η*_1,*i*_ is the neighbors-to-spot effect of gene *i. π* is the convolution parameter between spatially structured (*π* = 0) and unstructured (*π* = 1) noise.

| Protein | Genes | DIC | Parameter |  |  |  |  |
| --- | --- | --- | --- | --- | --- | --- | --- |
| | | | $\eta_{0,1}$ | $\eta_{0,2}$ | $\eta_{1,1}$ | $\pi$ | $\sigma^2$ |
| Podoplanin | 0 | 21625.401 | – | – | – | 0.005<br>(0.004) | 0.139<br>(0.005) |
|  | 1 | 21625.400 | 0.379<br>(0.052) | – | – | 0.006<br>(0.006) | 0.135<br>(0.005) |
|  | 2 | 21623.950 | 0.046<br>(0.094) | 0.624<br>(0.123) | – | 0.012<br>(0.010) | 0.121<br>(0.004) |
| I-A/I-E | 0 | 21620.370 | – | – | – | 0.013<br>(0.008) | 0.081<br>(0.003) |
|  | 1 | 21619.250 | 0.411<br>(0.052) | – | 0.019<br>(0.012) | 0.032<br>(0.026) | 0.076<br>(0.003) |
| CD117 | 0 | 12791.790 | – | – | – | 0.026<br>(0.016) | 0.191<br>(0.010) |
|  | 1 | 12790.650 | –0.116<br>(0.074) | – | –0.015<br>(0.017) | 0.032<br>(0.026) | 0.191<br>(0.010) |

## Notes

### Competing Interest Statement

The authors have declared no competing interest.

